# Identification of cardiac afterload dynamics from data

**DOI:** 10.1101/2021.03.31.437920

**Authors:** Henry Pigot, Jonas Hansson, Audrius Paskevicius, Qiuming Liao, Trygve Sjöberg, Stig Steen, Kristian Soltesz

## Abstract

The prospect of *ex vivo* functional evaluation of donor hearts is considered. Particularly, the dynamics of a synthetic cardiac afterload model are compared to those of normal physiology. A method for identification of continuous-time transfer functions from sampled data is developed and verified against results from the literature. The method relies on exact gradients and Hessians obtained through automatic differentiation. This also enables straightforward sensitivity analyses. Such analyses reveal that the 4-element Windkessel model is not practically identifiable from representative data while the 3-element model underfits the data. Pressure–volume (PV) loops are therefore suggested as an alternative for comparing afterload dynamics.

## 1. INTRODUCTION

### 1.1 Ex vivo heart evaluation

Today, donor hearts are routinely discarded due to uncertainty about their function. A method of functional evaluation of donor hearts therefore has the potential to increase the availability of heart transplantation.

Such *ex vivo* (outside of the body) functional evaluation requires a system that mimics vascular dynamics—the afterload—and can be controlled to emulate a broad range of recipient physiology and working conditions.

The afterload can be decoupled from heart dynamics by simultaneously measuring aortic flow and pressure, and relating them through an impedance model. Here, we delimit our focus to Windkessel models, being a class of linear time-invariant (LTI) arterial impedance models introduced by Otto Frank in the late 19^th^ century and commonly employed both academically and clinically.

We propose a method for identifying the Windkessel models from data, and perform sensitivity analyses to quantify uncertainty of the identified models. Three data sources are considered: previously published *in vivo* human data; porcine heart beating *in vivo*; porcine heart beating *ex vivo* against a synthetic afterload. The data and code used to generate the results are available on GitHub, see Pigot (2021).

This work focuses on technical aspects related to identifiability and parameter identification; details on the device and data generating experiments will be presented elsewhere.

### 1.2 The arterial Windkessel

Several variants of the Windkessel model have been proposed in the literature, see for instance Westerhof et al. (2009). Here we have employed the parallel 4-element Windkessel model, being one of the more general formulations. It can be expressed in terms of the circuit analogy shown in Fig. 1, that relates pressure (potential) *y_c_* to antegarde flow (current) ***u_c_*** through a dynamic impedance ***G_c_***, where we use subscript ***c*** to denote continuous time. The passive component parameters are described in table 1.

**Fig. 1.**
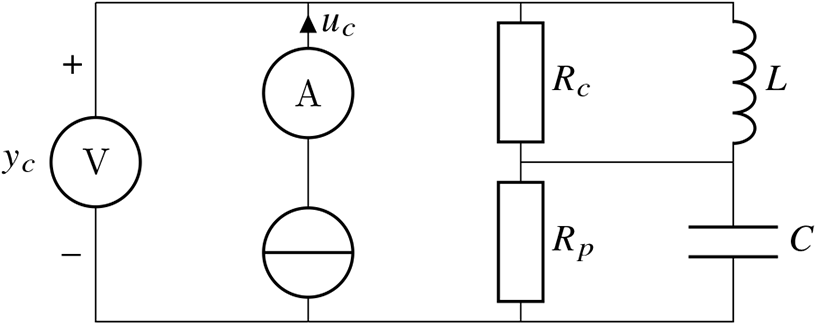
Circuit analogy of the parallel 4-element Windkessel model with driving current (flow) *u_c_*, corresponding voltage (pressure) *y_c_*, and parameters ***R_p_, C, R_c_, L.***

**Table 1.** Parameters of the Windkessel model shown in Fig. 1. Note that subscript *c* here denotes characteristic resistance (impedance) Westerhof et al. (2009), not to be confused with our use of subscript *c* to indicate continuous time.

| Parameter | Unit | Name |
| --- | --- | --- |
| $R_p$ | mmHg/(L/min) | Peripheral resistance |
| $C$ | L/mmHg | Compliance |
| $R_c$ | mmHg/(L/min) | Characteristic resistance |
| $L$ | mmHg min/(L/min) | Inertance |

Using the circuit analogy of Fig. 1 we next derive the expression for the transfer function ***G_c_*(*s*)**. In the Laplace domain the pressure (voltage) ***U*** across resistance ***R***, inertance (inductance) ***L*** and compliance (capacitance) ***C*** relate to the flow (current) ***I*** flowing though each element through ***RI*** = ***U***, ***sLI*** = ***U***, and ***I*** = ***sCU***, respectively. Denoting by ***p_c_*** the (pressure) potential between the resistances according to Fig. 1, Kirchhoff’s current law yields

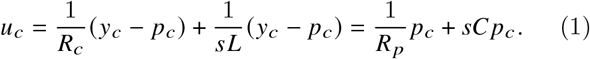

From (1) we can eliminate ***p_c_*** = ***R_p_***/(**1** + ***sCR_p_***)***u_c_*** to obtain the transfer function

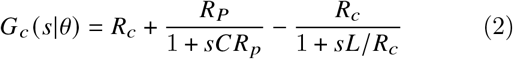

from ***u_c_*** to ***y_c_***, parameterized in ***θ*** = [***R_p_ C R_c_ L***]^⊤^ ≻ 0.

The Windkessel model (2) is hence a parallel interconnection between ***R_c_*** and two first-order systems. Note that the characteristic resistance ***R_c_*** only impedes accelerating flows, since for steady flows, the impedance of the inertance ***L*** is 0. This explains why the static gain of (2) is ***G***(0) = ***R_p_*** (and not, as some might assume ***R_c_*** + ***R_p_***). The 4-element Windkessel also comes in a less widespread series form, used in for example Gellner et al. (2020). In the series form, the inductance is connected in series with ***R_c_***, and its transfer function is obtained by replacing the last term in (2) by sL. The static gain is ***R_c_*** + ***R_p_***, but the transfer function is improper, as the term sL corresponds to an unfiltered differentiation of the input.

Introducing the state ***x_c_*** = [***x***_***c*1**_ ***x***_***c*2**_]^⊤^, defined through

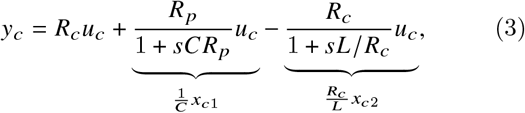

results in the state space form 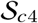 with matrices defined through

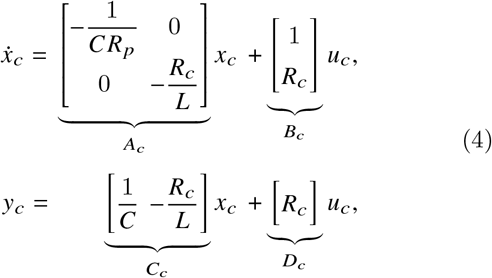

with *x*_*c*1_ being the volume (charge) in *C*. From (4) it is directly visible that the system has two poles corresponding to time constants ***T***_1_ = ***CR_p_*** and ***T***_2_ = ***L/R_c_***.

The 3-element and 2-element Windkessel models—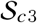 and 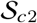—are commonly employed special cases of the 4-element version (4). In anticipation of Sec. 3, we note that ***L***/||*θ*|| **→** 1 results in 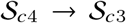, while either ***L*** → 0, ***R_c_*** → 0, or ***R_c_***/||*θ*|| → 1 result in 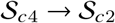.

## 2. METHOD

### 2.1 Experiments

The porcine data used in this work were recorded from two 65 kg Swedish pigs (*sus scrofa domesticus*). Large-animal experiments were needed to obtain a reliable characterization of the synthetic afterload module under consideration. All institutional and national guidelines for the care and use of laboratory animals were followed and approved by the appropriate institutional committees. The animals were treated in compliance with Directive 2010/63/EU, The European Parliament (2010). The study ran under ethics permission M174-15, issued by “Malmö/Lunds Djurförsöksetiska Nämnd” (local REB).

In both the *in vivo* and *ex vivo* experiments, a CardioMed CM4000 ultrasonic transit time flow meter (Medistim ASA, Oslo, Norway) was secured to the ascending aorta. Aortic pressure was measured using a Meritrans DTXPlus pressure transducer (Merit Medical, Singapore). In-house developed data acquisition hardware and software were used to log the signals at 200 Hz.

Different hearts were used for the *in vivo* and *ex vivo* data. The *in vivo* data reflects normal sinus rhythm in rest. The *ex vivo* data was recorded after 24 hours of cold non-ischemic perfusion, as described in Steen et al. (2016), beating against an actively controlled synthetic afterload akin to the device described by Gellner et al. (2020).

### 2.2 Model formulation

We relate the continuous-time flow ***u_c_*(*t*)** to the aortic pressure ***y_c_*(*t*)** through ***y_c_*** = ***g_c_*** * ***u_c_*** + ***ϵ_c_***, where ***g_c_*** is the impulse response of a an LTI system model with transfer function ***G_c_***, and ***ϵ_c_*** is the output error signal of the model, also referred to as the residual.

We do not have access to ***u_c_*** and ***y_c_*** directly, but only to corresponding measurement time series ***u, y***, each of ***n*** elements. The time series will here be considered equitemporarily sampled at a period of ***h***, although the proposed methodology is readily applicable also under irregular sampling schemes.

Applying the zero-order-hold **Ш**_***h***_ operator we thus obtain [***u, y, ϵ′, G***] = **Ш_*h*_** [***u_c_, y_c_, ϵ_c_, G_c_***] that relate through ***y*** = ***g*** * ***u*** + ***ϵ*** and discrete time state space realization 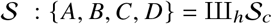

In order to evaluate the residual, *ϵ*, we simulate 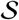 with *u* as input.The residual can be decomposed into one model-mismatch term and one noise term. The former typically arises from a candidate g under-modelling the data ***u, y***. The latter arises if y cannot be fully explained by *u*. This is for instance the case if *u* and *y* have been subjected to measurement noise. In this work we do not discern between the two contributors to *ϵ*.

To determine the initial state vector for this simulation, we could choose an arbitrary value, e.g. *x*_0_ = 0 and drive 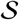 with a repeated stack [***u***^⊤^…***u***^⊤^]^⊤^ with sufficiently many repetitions of *u* for the transient caused by *x*_0_ to fade, and then discard all but the last *n* simulated output samples. An efficient and approximation-free alternative is to directly enforce the corresponding “periodic stationarity” condition *x*_0_ = ***x_n_***. Simulating the system forward in time over one cardiac cycle then gives

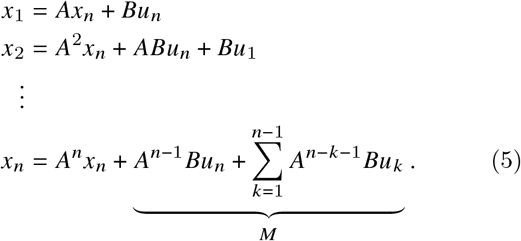

Using that ***x_n_*** = ***x*_0_** we solve (5) for

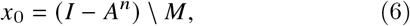

where, for numeric robustness, the sum in (5) is preferably computed as a convolution, rather than term-by-term.

### 2.3 Parameter identification

As stated in Sec. 2.2 we do not explicitly consider observation model (or other noise source) stochastics in this work, and therefore proceed with output error identification. The objective is hence to minimize the residual cost

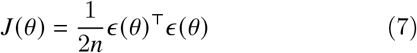

to identify

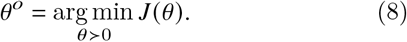

For a candidate ***θ*** we can evaluate 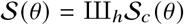, and obtain *x*_0_ using (5). Driving 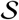 with ***u***, we then obtain the output 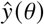 and residual 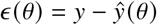.

The solution *θ*^o^ of (8) will also minimize 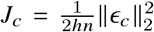, under the assumption that ***u_c_*** is piece-wise constant between the samples in ***u***, and that the signals ***u_c_*** and ***y_c_*** have been subjected to analog filtering effectively removing spectral power above the Nyquist frequency **(2*h*)^-1^**. The latter holds true for the data considered here. The former constitutes a valid approximation as long as 1/*h* is large compared to the magnitudes of the poles of ***G***. Validity of this approximation has been ensured through sampling at a high frequency compared to the cardiac cycle dynamics.

The optimization problem (8) is generally non-convex in *θ*. We therefore employ a multiple initialization procedure. Being the static gain of the system, ***R_p_*** is initialized with the mean output (pressure) divided by mean input (flow). The initial values of the remaining parameters are drawn from a multivariate uniform distribution, and we retrospectively verify that this distribution covers the identified parameters *θ*^o^.

For each initialization point, Newton’s method is then used to obtain a cost minimizer candidate, and the one of them that minimizes ***J*** gets denoted *θ*^o^.

A dual number implementation (see Revels et al. (2016)) of the zero-order-hold operator III_*h*_ enabled *exact* (down to machine precision) forward-mode automatic differentiation and thus evaluation of the gradient **∇*J*(*θ*)** and Hessian **∇**^2^*J*(*θ*) required in each Newton iteration. This enables us to identify the continuous-time model parameter *θ* directly, without the need of finite difference, or other, approximations. It also means that we can obtain an *exact* evaluation of the Hessian ***H* = ∇^2^*J*(*θ*^o^)** of the cost ***J*** with respect to the parameter *θ*, evaluated at the optimum *θ*^o^.

### 2.4 Persistence of excitation

A classical result for identification of LTI systems is that the order of persistence of excitation (PE) of an input sequence *u* determines whether *u* is sufficiently informative to distinguish parameter candidates *θ* and *θ*′ given some model class, or structure, *G*. Particularly, PE of order at least *m* is required for *u* to be sufficiently informative to distinguish any pair of transfer functions of *m* parameters. See for example Mareels et al. (1987) for further detail.

The PE degree can be determined in several (equivalent) ways, one being through a rank condition on the expectation of the auto-correlation matrix of *u*:

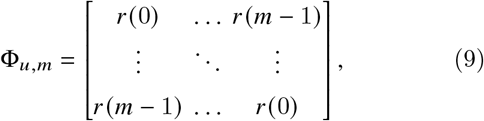

where ***r*(*τ*)** is expectation of the auto-correlation of *u*, with respect to some stochastic observation model relating *u_c_* to *u*. In absence of stationary (additive) noise model, it is customary to assume that the observation *u_c_*(*kh*) is an unbiased and consistent estimate of ***u*(*k*)** and use the observed auto-correlation, defined through (9) with

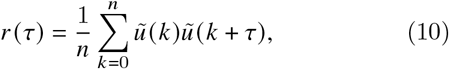

where 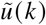 is typically defined as ***u*(*k*)** for 1 ≤ ***k*** ≤ ***n***, and 0 otherwise. Since we are dealing with signals of periodic nature here, we will instead use the periodic expansion 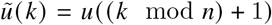. We can efficiently evaluate the corresponding circular sample auto-correlation sequence

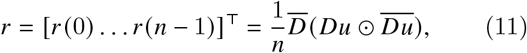

where ***D*** is the *n* × *n* DFT matrix, 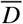 its conjugate, and ⊙ denotes element-wise matrix-matrix multiplication.

The rank condition states that in order for ***u*** to be PE of order at least ***m***, the corresponding **Φ_u_** needs to be positive definite. A major concern here is that any ***u*** can be turned into a signal of arbitrarily high PE order by adding a signal of independent samples 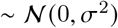, even with arbitrarily small ***σ*** > 0. From a practical perspective it is therefore more relevant to consider the spectrum of **Φ_*u*_**.

### 2.5 Sensitvity analysis

Next, we investigate how much model fit, expressed in terms of the cost ***J***, deteriorates if the optimal parameter *θ*^o^ is subject to an additive perturbation with magnitude ||*δ*||_2_. The Taylor series expansion of the perturbed cost is

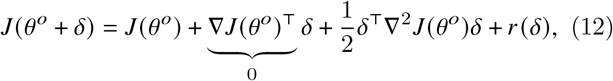

where the residual *r*(*δ*) is a linear combination of monomials in the components of *δ*, each with degree at least 3. If ||*δ*||_2_ is small, then the contribution of *r*(*δ*) to ***J*** is small. In that case the cost increment is well approximated by

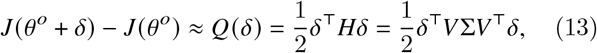

where the Hessian ***H* = ∇^2^*J*(*θ*°)** is a symmetric real matrix and therefore has a singular value decomposition according to (13). The singular vectors make up the columns of the unitary matrix *V* = [*v*_1_…*v_m_*] and we can write the quadratic form as

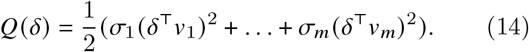

For a fixed ||*δ*||_2_, (14) is minimized (maximized) when *δ* is parallel to the singular vector *v_k_* corresponding to the smallest (largest) singular value *σ_k_*. This reveals in what direction a small move away from *θ*° contributes least (most) to increase ***J***. Further, the fraction between the largest and smallest singular value, being the condition number of *H*, reveals the relative change in cost when moving a small (infinitesimal) distance ||*δ*||_2_ in the least and most sensitive directions, respectively. A large condition number is therefore an indicator of over-parametrization with respect to the experimental data.

## 3. RESULTS

### 3.1 Identified models

The identification method was first validated against a previously published human data set—henceforth refered to as the *human* data—by digitizing the waveforms in Figure 4 of Stergiopulos et al. (1999) using Webplotdigitizer, Rohatgi (2020). Numerical values are reported in table 2; time domain model outputs and corresponding pressure– volume (PV) loops in Fig. 2; output error residuals in Fig. 5. As seen in Fig. 2, the parameter values identified using the method of Sec. 2 reproduce the appearance of the results in Stergiopulos et al. (1999). For reasons explained above in Sec. 2.5, 2- and 3-element Windkessel models were also identified.

**Fig. 2.**
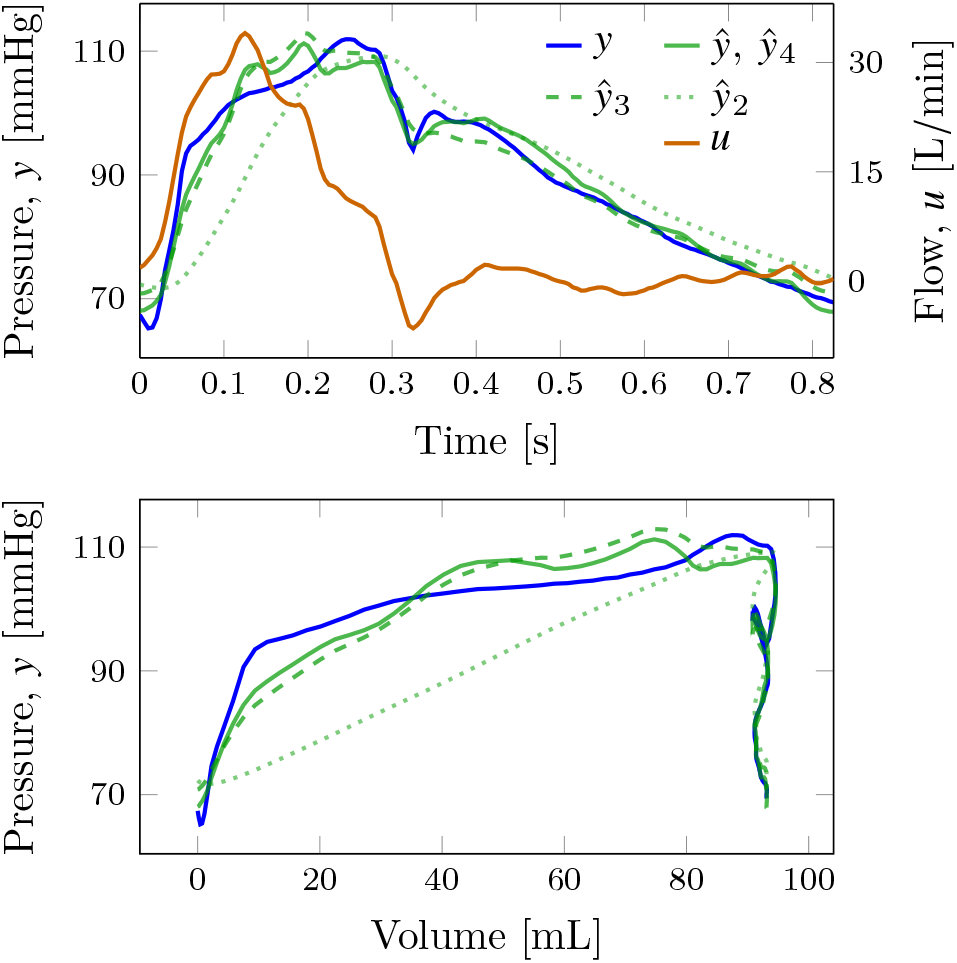
Data set *u, y* and reported 4-element Windkessel model output 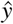 digitized from Fig. 4 of Stergiopulos et al. (1999); model output 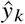 of the herein identified *k*-element Windkessel models. Top pane shows time domain model fit; bottom pane shows PV loop.

**Table 2.** Identified parameters of 4-, 3-, and 2-element Windkessel models together with mean squared error (MSE) [mmHg], directly proportional to ***J*(*θ*°)** of (7). The last column shows the conditioning of ***H* = ∇^2^*J*(*θ*)** at *θ* = *θ* °.

| | $R_p$ | $C$ | $R_c$ | $L$ | MSE | cond($H$) |
| --- | --- | --- | --- | --- | --- | --- |
| Human | 13.6 | 0.0743 | 0.952 | 0.0952 | 5.88 | 3.18e3 |
|  | 13.0 | 0.108 | 0.582 |  | 8.48 | 3.65e2 |
|  | 13.6 | 0.0996 |  |  | 48.2 | 3.07e2 |
| *in vivo* | 62.9 | 0.0855 | 1.58 | 8.52 | 30.9 | 1.20e7 |
|  | 61.3 | 0.0877 | 1.58 |  | 30.9 | 1.57e3 |
|  | 62.9 | 0.0853 |  |  | 77.1 | 1.37e3 |
| *ex vivo* | 148 | 0.345 | 0.615 | 31.3 | 7.16 | 2.76e9 |
|  | 148 | 0.346 | 0.615 |  | 7.16 | 1.00e2 |
|  | 147 | 0.353 |  |  | 23.3 | 5.56e1 |

Moving on to the experimental data, identified parameter values, mean squared error (MSE) of the time domain model fit, being directly proportional to ***J*(*θ*°)**, and the condition number of the Hessian ***H* = ∇^2^*J*(*θ*°)** are listed in table 2. Time domain model outputs and PV loops are shown in Fig. 3 and Fig. 4; output error residuals are shown in Fig. 5.

**Fig. 3.**
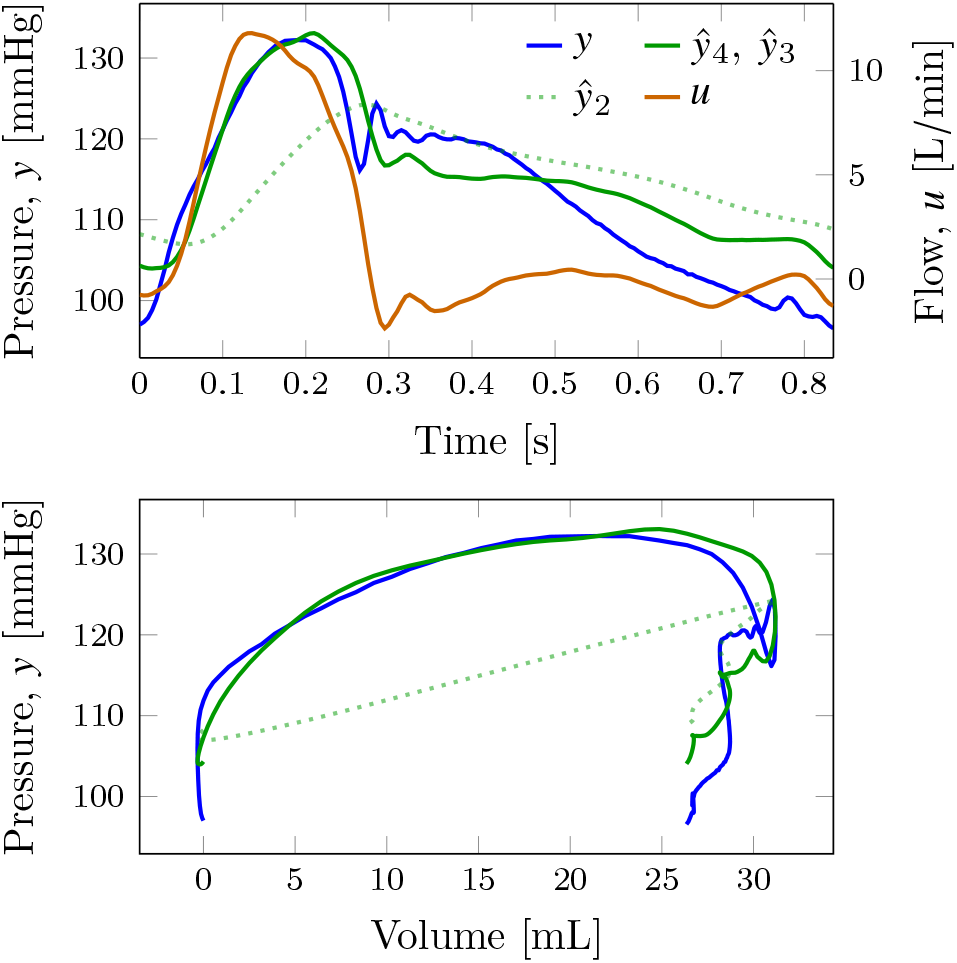
*In vivo* data set *u, y* together with outputs 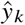 of identified *k*-element Windkessel models.

**Fig. 4.**
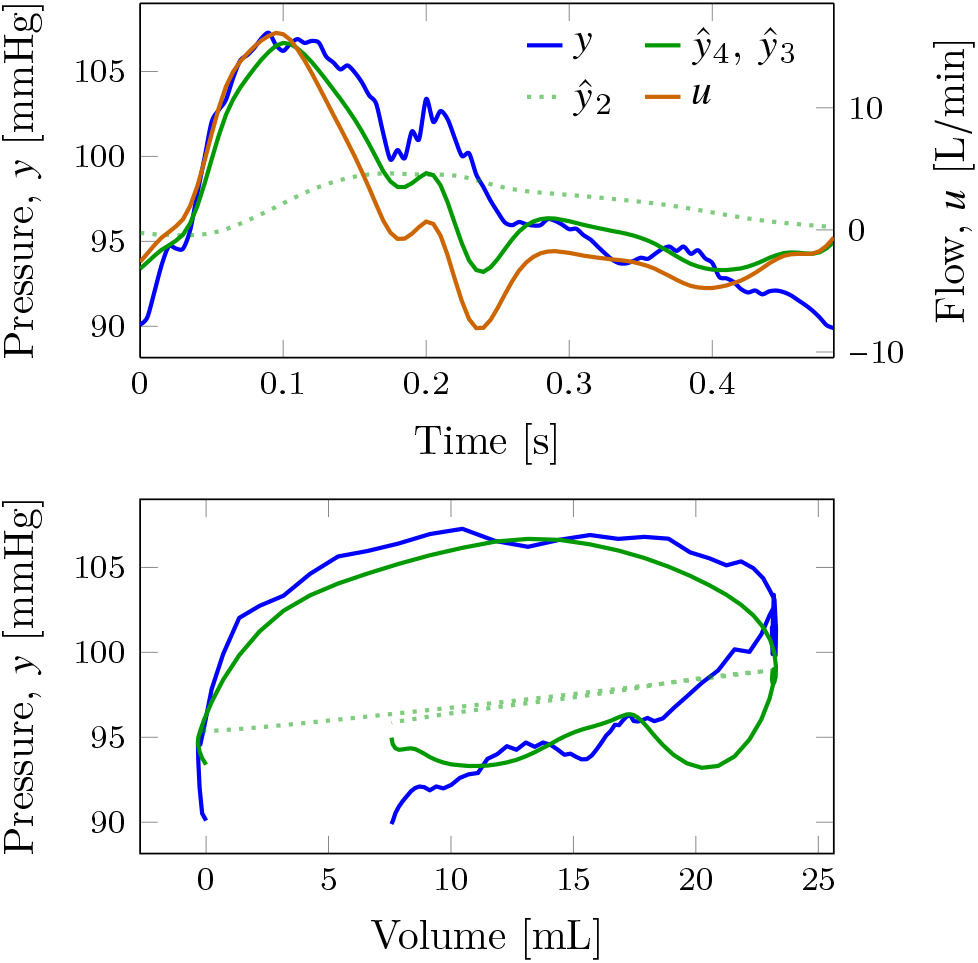
*Ex vivo* data set *u, y* together with outputs 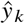 of identified *k*-element Windkessel models.

**Fig. 5.**
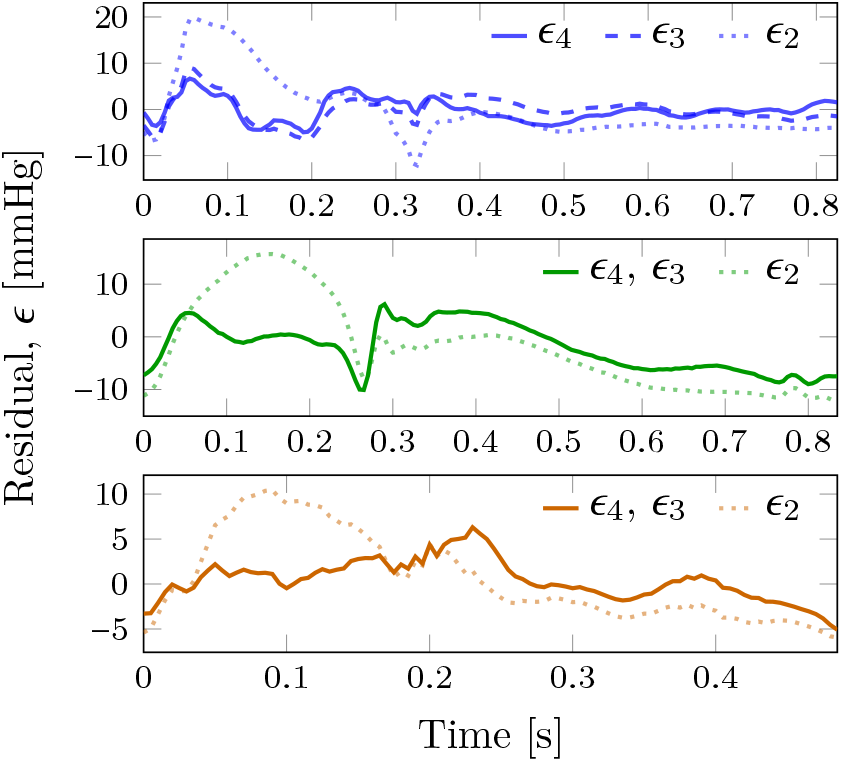
Output error residuals *ϵ_k_*, of *k*-element Windkessel models identified for the human (top); *in vivo* experiment (middle); *ex vivo* experiment (bottom).

Bode plots of the identified models are shown in Fig. 6. These indicate that for the experimental data an almost perfect pole-zero cancellation takes place in the 4-element models, as can be verified by inserting the parameter values of table 2 into (2). This also explains why corresponding time-domain plots are visually indistinguishable.

**Fig. 6.**
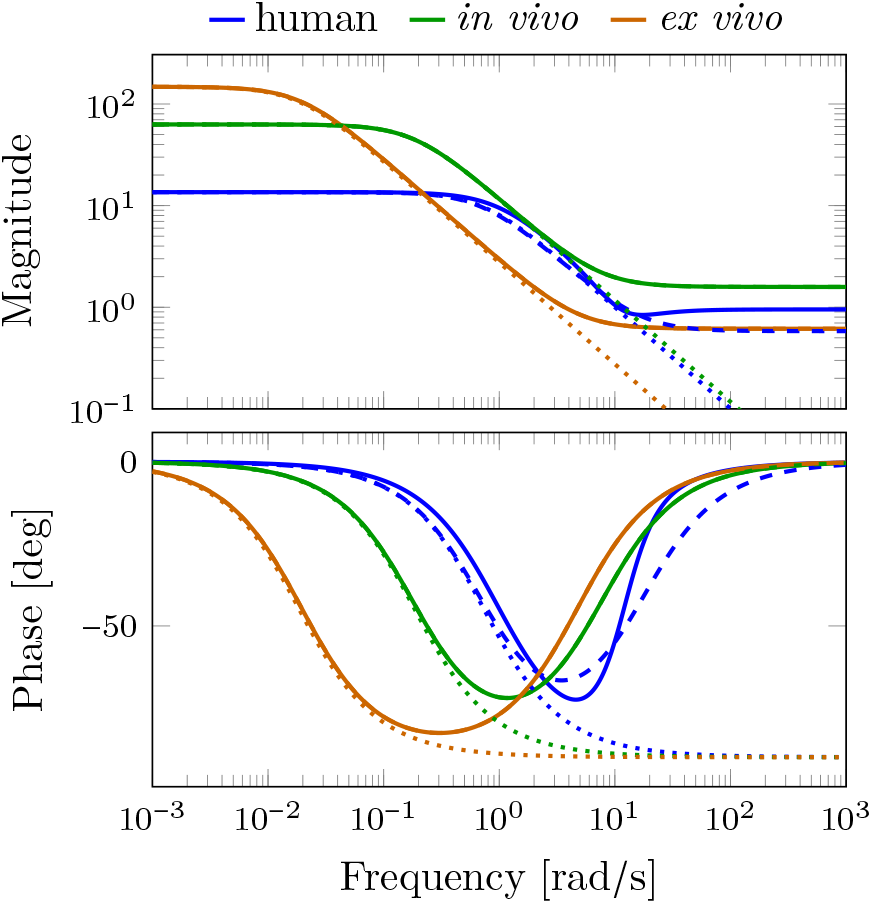
Bode plots of the identified 4-element (solid), 3-element (dashed), and 2-element (dotted) Windkessel models from the human (blue), *in vivo* (green) and *ex vivo* (bronze) data sets.

### 3.2 Persistence of excitation

The 10 largest singular values of the auto-correlation matrices **Φ_*u*_** for the inputs of the human, *in vivo*, and *ex vivo* data sets are shown in Fig. 7. According to Sec. 2.4, the figure indicates that identification of 2 to 4 parameters may be feasible, which prompts further consideration of parameter sensitivity in the fitted models. Since the input is generated by the heart, it cannot be arbitrarily changed to increase excitation and elucidate more parameters.

**Fig. 7.**
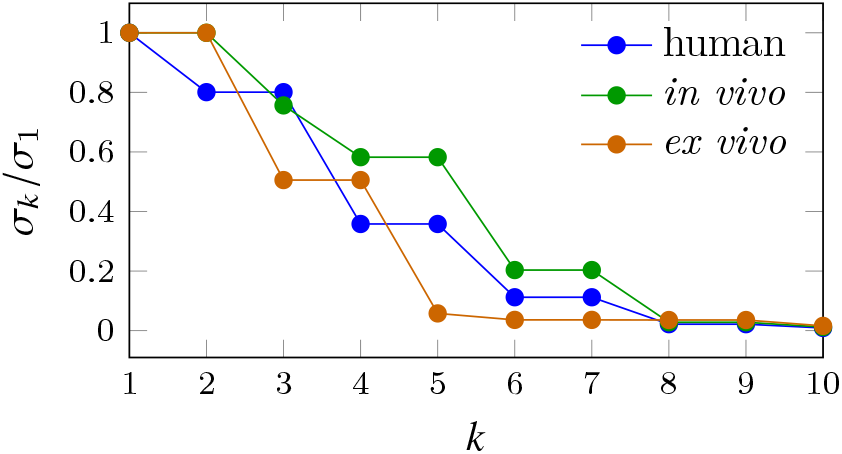
The 10 largest singular values *σ*_1_ ≥ … ≥ *σ*_10_ of **Φ_*u*_**, normalized by *σ*_1_. The spectral densities **Φ_*u*_** were computed using ***u*** from human (blue); *in vivo* data set (green); *ex vivo* data set (bronze).

One can note that two parameters (degrees of freedom) are necessary to reproduce arbitrary diastolic and systolic pressure levels. For the 4-element Windkessel model 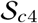, this leaves two parameters to influence the shape of the model output; one “shape” parameter for 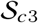; none for 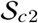.

### 3.3 Parameter sensitivities

The singular value decompositions *H* = *V*Σ*V*^⊤^ of the Hessian ***H* = ∇^2^*J*(*θ*)**, evaluated at the identified parameter *θ* = *θ*° are shown in table 3. The last column of the *V* matrices indicate that the least certain direction in parameter space coincides with ***R_c_*** for the human data. The *in vivo* and *ex vivo* data instead result in uncertain estimates of *L*. Recalling from Sec. 1.2 how limit cases of ***L*** and ***R_c_*** correspond to the 2- and 3-element Windkessel structures, it is not surprising—given the indicated sensitivities—that the 3-element models explain the experimental data almost identically well, while the 2-element counterparts fail to do so based on lacking degrees of freedom, as mentioned in Sec. 3.2.

**Table 3.**
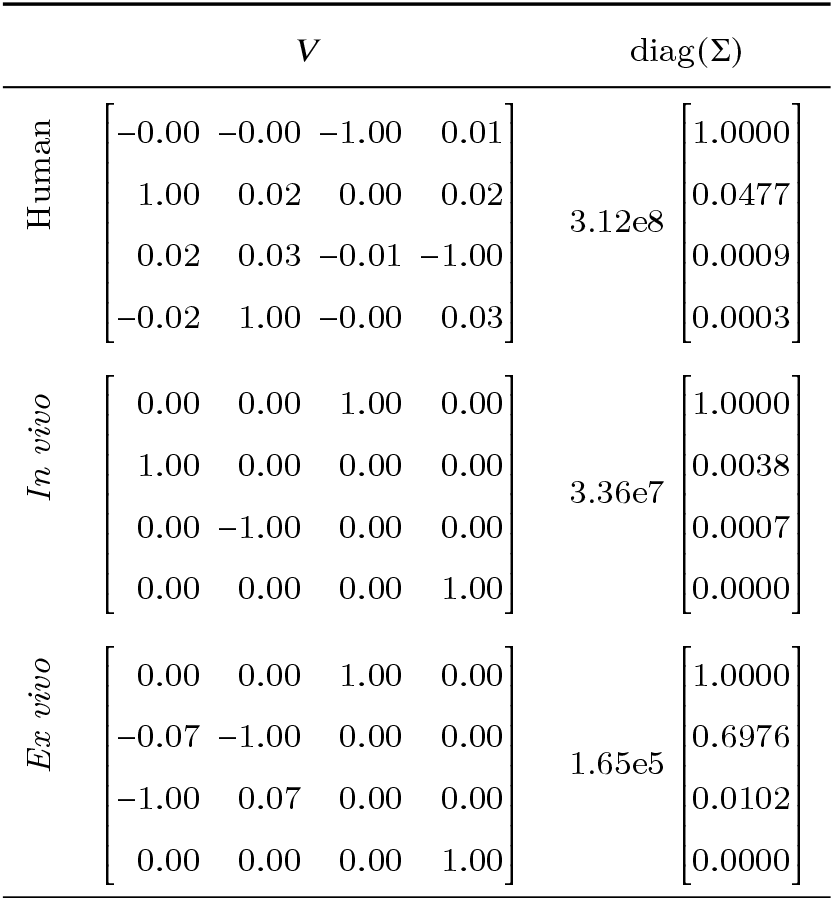
Singular value decomposition ***H* = *V*Σ*V*^⊤^** of the Hessian ***H* = ∇^2^*J*(*θ*)**.

## 4. DISCUSSION

A method for identification of continuous time dynamics from time series data has been proposed with the objective of comparing the dynamics of a synthetic afterload with those of normal physiology.

Upon validation against previously published results from Stergiopulos et al. (1999), the method was therefore applied to two porcine data sets: one collected *in vivo*, featuring normal physiology; one collected *ex vivo*, with the heart working against a synthetic afterload, as may be used in functional evaluation of donor hearts prior to possible implantation.

We can note that model fit of the human data is slightly better than for our experimental data. This is presumably caused by how our measurement setup was devised, and we are planing to conduct similar but refined measurements to investigate this. Nonetheless, the output error residuals from all three data sets, shown in Fig. 5, suggest that the Windkessel structure under-models representative pressure–volume data. At the same time, the persistence of excitation analysis of Sec. 2.4 and local sensitivity analysis of Sec. 2.5 suggest that the inputs ***u*** do not support identification of substantially more complex LTI model structures than the 3-element Windkessel.

## 5. CONCLUSION

The well-established 4-element parallel Windkessel model is not reliably identifiable from representative pressure– volume time series data. Particularly, the peripheral resistance ***R_p_***, being the static gain of the model, is reliably identifiable, while inertance parameter ***L*** is unidentifiable. For the sake of comparing afterload impedance dynamics, it is therefore advisable to look directly at the PV loop representation, rather than comparing Windkessel model parameters.

## 6. ACKNOWLEDGMENTS

This work was partially funded by the Swedish Research Council (grant 2017-04989) and the Wallenberg AI, Autonomous Systems and Software Program (WASP) funded by the Knut and Alice Wallenberg Foundation. The authors from the Department of Automatic Control are members of the ELLIIT Strategic Research Area at Lund University.

